# Increased microbial diversity and decreased prevalence of common pathogens in the gut microbiomes of wild turkeys compared to domestic turkeys

**DOI:** 10.1101/2021.07.16.452759

**Authors:** Julia Craft, Hyrum Edington, Nicholas D. Christman, John Chaston, David Erickson, Eric Wilson

## Abstract

Turkeys (*Meleagris gallopavo*) provide a globally important source of protein and constitute the second most important source of poultry meat in the world. Bacterial diseases are common in commercial poultry production causing significant production losses for farmers. Due to the increasingly recognized problems associated with large-scale/indiscriminant antibiotic use in agricultural settings, poultry producers need alternative methods to control common bacterial pathogens. In this study we compared the cecal microbiota of wild and domestic turkeys, hypothesizing that environmental pressures faced by wild birds may select for a disease-resistant microbial community. Sequence analysis of 16S rRNA genes amplified from cecal samples indicate that free-roaming wild turkeys carry a rich and variable microbiota compared to domestic turkeys raised on large-scale poultry farms. Wild turkeys also had very low levels of *Staphylococcus, Salmonella* and *E. coli* when compared to domestic turkeys. *E. coli* strains isolated from wild or domestic turkey cecal samples also belong to distinct phylogenetic backgrounds and differ in their propensity to carry virulence genes. *E. coli* strains isolated from factory-raised turkeys were far more likely to carry genes for capsule (*kpsII*, *kpsIII*) or siderophore (*iroN*, *fyuA*) synthesis than those isolated from wild turkeys. These results suggest that the microbiota of wild turkeys may provide colonization resistance against common poultry pathogens.

**Importance:** Due to the increasingly recognized problems associated with antibiotic use in agricultural settings, poultry producers need alternative methods to control common bacterial pathogens. In this study we compare the microbiota of wild and domestic turkeys. Results suggest that free ranging wild turkeys carry a distinct microbiome when compared to farm raised turkeys. The microbiome of wild birds contains very low levels of poultry pathogens compared to farm raised birds. The microbiomes of wild turkeys may be used to guide development of new ways to control disease in large scale poultry production.

## Introduction

Turkeys (*Meleagris gallopavo*) evolved approximately 11 million years ago and are one of the first birds domesticated in the Americas (1–3). Although domesticated thousands of years ago, turkeys have remained generally very similar to their wild relatives until relatively recently (4, 5). In the past ~70 years, intensive selective breeding of turkeys has resulted in dramatic changes in commercially raised birds compared to their wild relatives, leading to a genome that is much less diverse than many other agricultural species (4). These genetic changes as well as advancements in production practices have resulted in domestic birds maturing much more quickly and reaching three times the body mass of wild birds at maturity (6). Domestic turkeys are now the second most important source of poultry in the world, with the USA producing ~250,000,000 turkeys and ~7,000,000,000 pounds of turkey meat in 2019 (7).

Relatively few studies have been published comparing the microbiomes of wild animals and their domesticated kin. However, the limited literature on this topic has overwhelmingly shown that the microbiome of captive and wild animals varies dramatically (8–15). The observed differences in microbial communities between wild and captive animals has led for calls for more research on the microbiomes of additional wild animals (16, 17).

The gut microbiome of poultry is known to contribute to efficient growth as well as bird health (11, 18–21). The microbiome of commercially raised poultry is undoubtedly influenced by production practices such as crowded conditions, diet, and antibiotic use. Several studies have characterized the gut microbiomes of domestic turkeys in a variety of experimental and agricultural settings (20, 22–25); however very few studies have focused on the microbiomes of wild turkeys (11).

In an effort to better characterize potential effects of gut microbiota on turkey health and disease, we compared the cecal microbiota from factory-raised domestic, free-ranging domestic and free-ranging wild turkeys. Sequencing of the V4 region of 16S DNA was used to determine the abundance of multiple taxa in the ceca of individual birds within each group. Additional experiments were designed to determine the prevalence of bacterial taxa which are common pathogens of commercially raised turkeys. These studies indicate that beta diversity within the microbiota is significantly different between factory-raised domestic turkeys, free-ranging domestic turkeys and free-ranging wild turkeys. Several common pathogens associated with commercial poultry production (*E. coli*, *Salmonella and Staphylococcus sp)*, were infrequent or absent in the cecal microbiota of free-ranging wild turkeys. *E. coli* strains found in wild turkeys were found to be genetically diverse and carry fewer virulence associated genes than strains found in factory-raised birds.

## Materials and Methods

### Definition of turkey groups used in this study

The term “wild turkey” can mean both a strain of turkey, as well as the lack of domestication. In this study, we define “wild turkey” as a population of self-sustaining, wild, free-ranging birds. All wild turkeys sampled in this work were of the Rio Grande subspecies (*Meleagris gallopavo intermedia*), that have ranged freely for generations in the mountains of North Central Utah, USA. Birds described as “free-range domestic turkeys” in this study are domesticated turkeys ranging freely outdoors. All free-range domestic turkeys in this study were from hobby farms where they were allowed to forage freely outdoors both summer and winter. The diet of all domestic free-range turkeys was supplemented with commercial poultry food by their owners. The term “factory-raised domestic turkey” refers to turkeys raised in commercial turkey production facilities. All turkeys in this group were of the *Broad Breasted White* variety. Although these factory-raised birds may fit the legal definition of “free-range” by virtue of their caging conditions, they were not considered “free-ranging” for the purposes of this study.

### Collection of cecal samples

Some birds, including turkeys and chickens, produce two distinctly different types of feces. Cecal drops are a type of feces that the bird periodically excretes directly from the intestinal cecum (26). Previous work has demonstrated that the ceca contains the greatest microbial diversity found in the intestinal tract of poultry (27, 28). Additionally, the microbiota found in cecal drops is highly reflective of the microbiota found in cecal contents collected following sacrifice of the bird (29). The collection of cecal drops, which are easily distinguishable from normal feces, enables a simple, noninvasive method of obtaining a clear view of the cecal microbiota and eliminates the need to sacrifice (or even come in contact with) study animals.

In this study all samples were of cecal origin. Cecal drops from wild and free-ranging domestic turkeys were collected during winter months following snowstorms. Sample collection immediately following snowstorms ensured that only fresh samples were collected and the sample remained relatively uncontaminated by bacteria from the soil or other environmental sources. Cecal contents from one flock of factory-raised turkeys were collected from a turkey processing facility post mortem. Cecal drops from a second commercially raised flock were collected from the floor of the production facility. Sampling sites, bird age and other details of sample origin are listed in Supplementary Data S1

### DNA preparation

Following sample collection, all cecal contents were kept frozen until DNA isolation. DNA used for V4 sequencing was extracted from each sample using the Zymo *Quick*-DNA Fecal/Soil Microbe 96 Kit (Zymo D6011) according to the manufacturer’s instructions, including a bead homogenization step using a 2010 Geno/Grinder (Spex, Metuchen, NJ) at 1750 RPM for 10 min. DNA was prepared for 16S rRNA gene V4 region sequencing based on an established protocol with minor deviations (30). First, the V4 region of the 16S rRNA gene was amplified individually from each sample with the AccuPrime Pfx Enzyme (ThermoFisher Scientific, Waltham, MA, USA) in 20 μl volumes using a subset of the exact primer sequences described previously (30). PCR amplicons were normalized using the SequalPrep Normalization kit (Applied Biosystems, Waltham, MA, United States), pooled in groups of 96 reactions, and fragments in the range of 250-450 bp were purified using a BluePippin (Sage Science, Beverly, MA) selection step. Equimolar normalization of each pool and sequencing was performed at the BYU DNA Sequencing center on a partial 2 × 250 lane (v2) of a HiSeq 2500 (Illumina, Inc., San Diego, CA). Laser complexity was assured by including at least 10% of each lane with shotgun sequencing libraries for other bacterial genomes. Sequences were deposited to the National Center for Biotechnological Information Short Read Archive as (accession forthcoming).

### Sequence Analysis

Sample reads were demultiplexed on the Illumina platform and analyzed using QIIME2 (31, 32) and R. briefly, reads were trimmed to maximize quality scores of each nucleotide position. DADA2 (33) was used to denoise, dereplicate, and call amplicon sequence variants (ASVs), taxonomy was assigned to the ASVs using the GreenGenes classifier 13_8_99 (34). ASV tables were filtered to 13,000 reads per sample and differences between groups were determined by PERMANOVA (35) of weighted and unweighted Unifrac distances (36, 37). To permit calculating Unifrac distances we built a phylogenetic tree with fasttree2 (38) based on mafft alignment (39). Differences in OTU abundances between samples were performed using ANCOM (40). Abundances of individual OUTs were manually analyzed based on the taxonomic assignments, which were assigned to OTUs using the QIIME2 q2-feature-classifier (41). Alpha diversity metrics were defined using QIIME2 and differences in alpha diversity metrics between sampling locations was determined by a Kruskal-Wallis test.

### Determination of relative E. coli DNA levels in cecal samples

A qPCR based assay was designed to estimate the relative abundance of *E. coli* DNA in each cecal sample, based on detection of the *ybbW* gene, which is found exclusively in *E. coli* (42). Primer probe sets and reaction conditions can be found in Supplemental Data (Figure S2). The efficiency and reproducibility of amplification was verified by generating a standard curve using doubling dilutions of positive control DNA. Negative controls consisted of reaction mixtures with DNA elution buffer rather than DNA. Each sample was tested in duplicate. Purified DNA from pooled *E. coli* strains was used as a positive control.

### Isolation and Genotyping of *E. coli* strains

*E. coli* present in cecal samples were isolated by homogenizing a portion of the sample in sterile PBS and plating on MacConkey agar, followed by growth at 37 °C for 24 h. Colonies with characteristic *E. coli* morphology were then restreaked and verified as *E. coli* by PCR targeting the *ybbW* gene. Total DNA was isolated from individual colonies using mini-genomic DNA kit for blood and cultured cells (IBI Scientific). Putative *E. coli* strains were assigned to phylo-groups using the Clermont quadriplex assay, with additional PCR tests to distinguish group C or group E when warranted, as previously described (43). Presence or absence of genes associated with virulence of avian pathogenic *E. coli* (*iutA*, *iss*, *iroN, fyuA, kpsMTII, kpsMTIII*) was determined by PCR, using primers contained in Supplemental Data S2.

### Detection of *Salmonella* DNA in cecal samples

Presence or absence of *Salmonella* sp. in cecal sample DNA was determined using a semi-quantitative PCR assay based on detection of the *invA* gene, which has previously been demonstrated to specifically detect most *Salmonella* strains (44). Primer sequences and PCR conditions are outlined in Supplemental Data S2.

## Results

To explore potential differences in the microbiota of wild vs domestic turkeys, a 16S rRNA gene survey of the cecal microbiota was performed. A total of 4,070,891 bacterial reads were obtained, with an average of 53,564 reads per sample and 3,069 ASVs. We performed principal coordinate analysis (PCoA) and PERMANOVA of weighted Unifrac distances to compare the microbiota composition of different flocks of turkeys. At a 13,000-read subsampling depth PERMANOVA of Unifrac distances revealed significant differences in the microbiota of the samples within provenance and flock (Table 1, Supplementary Data S3). The clustering of samples on PCoA ordinations visually depicted these statistical differences, where Principal coordinates 1 and 2 separated the samples into three general groups that matched the provenance of the samples when analyzed by both weighted and unweighted Unifrac distance (Figure 1, Table 1). Follow up weighted and unweighted Unifrac analyses confirmed there were flock-specific effects when each provenance was analyzed separately, except for birds from factory-raised flocks, analyzed by weighted Unifrac (Table 1). The finding that all flocks differed in beta diversity, except those raised in commercial production facilities, is likely a reflection of the highly standardized nature of commercial poultry production.

**Table 1.**
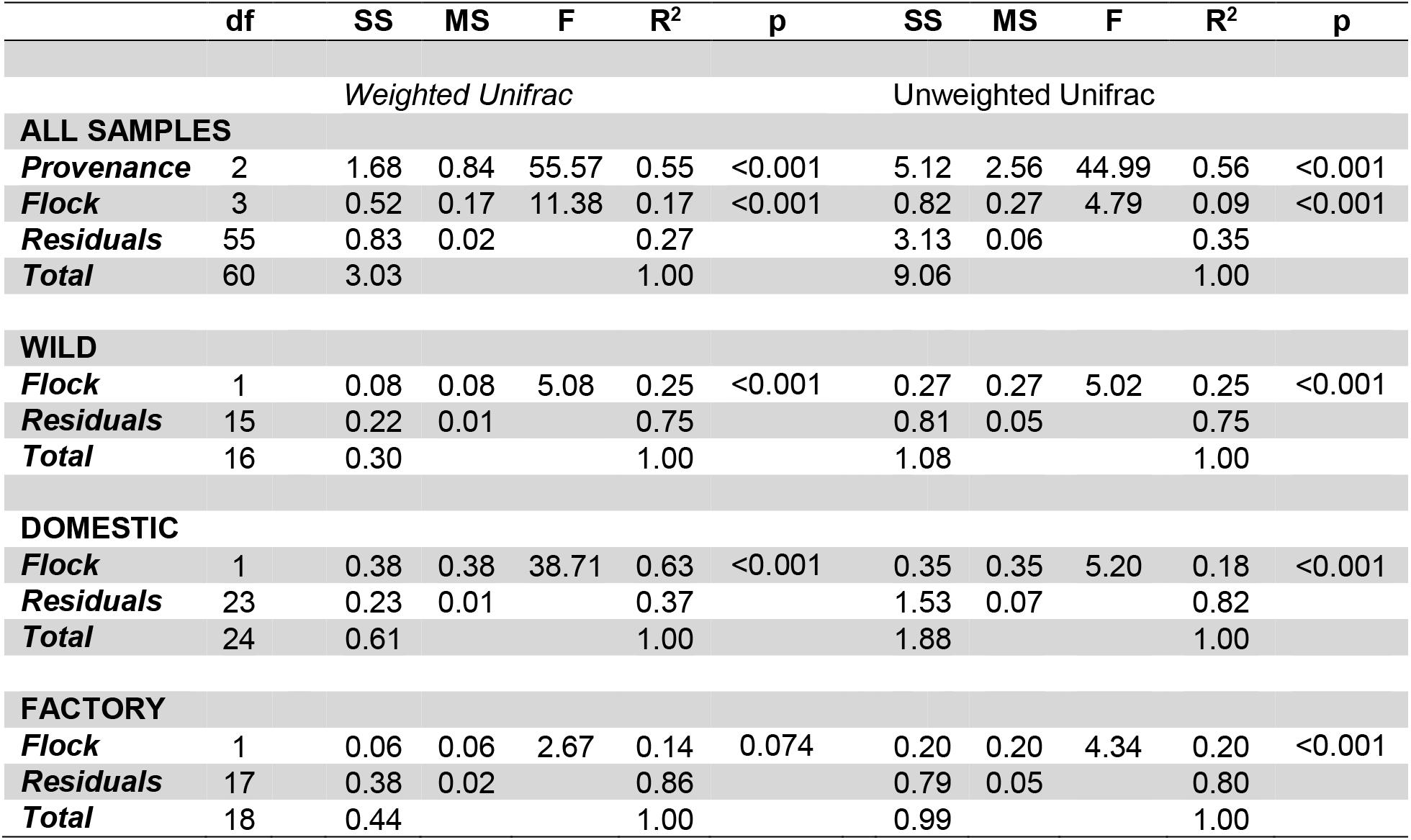
PERMANOVA tables for different groups of samples. Df = degrees of freedom, SS = sum of squares, MS = mean of squares, F = F statistics, R^2^ = R^2^ value. P = p-value.

**Figure 1.**
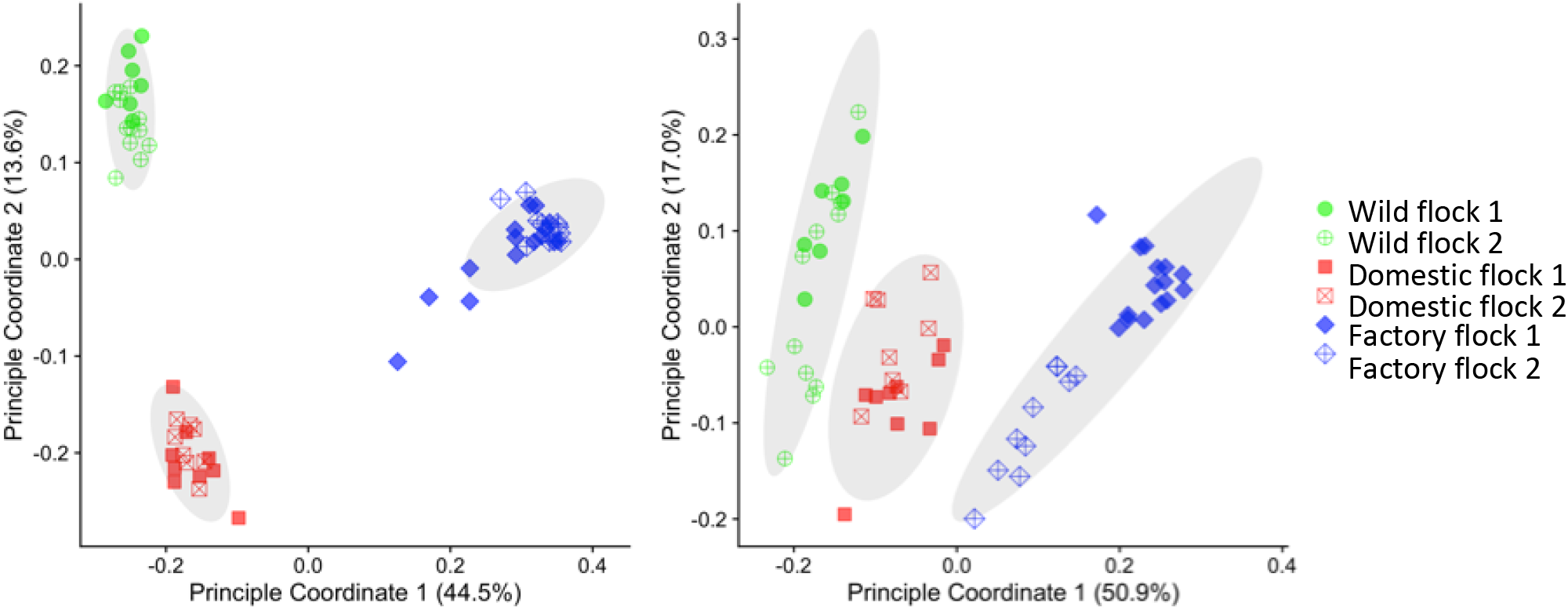
Principal Component Analysis demonstrates the cecal microbiota of turkeys’ cluster according to bird provenance. Weighted (A) and unweighted (B) Unifrac distance plots of the different flocks, colored according to the animals’ provenance.

We also evaluated the variation in the microbiota composition of the different flocks. At the order level there were significant differences in the numerical density of the most abundant bacterial taxa (Figure 2A). For example, in the factory-raised birds, *Clostridiales* was the most abundant taxon (71.7% +/− 3.3%), much more than in free-ranging domestic turkeys (33.8% +/− 1.8%) or wild turkeys (18.3% +/− 0.7%). The lower *Clostridiales* read counts in the free-ranging domestic and wild turkeys were largely offset by relative increases in *Bacteroidales* and *Coriobacteriales*. The abundances of these reads were all significantly different between provenances by ANCOM (Supplementary Data S4). However, despite these differences in abundance of different taxa, variation in alpha diversity between flocks was not related to the flocks’ provenance (Table 2). Therefore, key differences in numerical composition at high taxonomic levels did not necessarily reflect low-level differences in diversity.

**Figure 2A.**
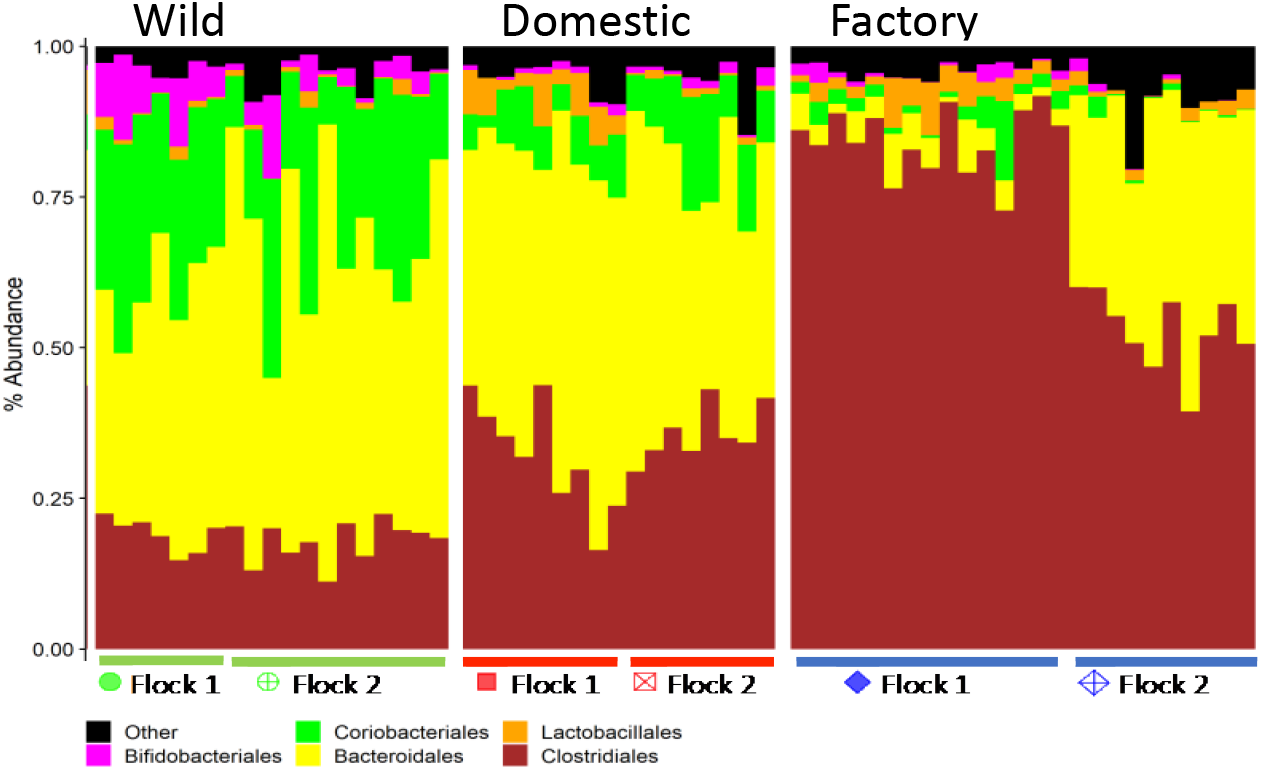
Taxa abundance differs widely between wild and factory raised domestic turkeys. Taxon plot of flocks, grouped by provenance. Order level assignments above 2% total relative abundance are shown individually.

**Table 2.** Alpha diversity metrics on a per-flock basis as mean +/− s.e.m.. Different letters next to the s.e.m. represent significant differences between flocks for each metric and were determined by a Kruskal-Wallis test

| Flock | Shannon | Pielou's Evenness | Faith's PD | Observed ASVs |
| --- | --- | --- | --- | --- |
| Wild flock 1 | 6.03 +/- 0.09 <sup>ab</sup> | 0.753 +/- 0.009 <sup>ab</sup> | 17.2 +/- 0.9 <sup>a</sup> | 260 +/- 13 <sup>ab</sup> |
| Wild flock 2 | 6.43 +/- 0.10 <sup>cd</sup> | 0.77 +/- 0.008 <sup>ac</sup> | 21.2 +/- 0.7 <sup>b</sup> | 330 +/- 18 <sup>c</sup> |
| Domestic flock 1 | 6.61 +/- 0.12 <sup>ac</sup> | 0.785 +/- 0.010 <sup>a</sup> | 21.1 +/- 0.9 <sup>b</sup> | 347 +/- 21 <sup>ad</sup> |
| Domestic flock 2 | 6.96 +/- 0.03 <sup>ac</sup> | 0.81 +/- 0.007 <sup>a</sup> | 22.4 +/- 0.8 <sup>a</sup> | 402 +/- 20 <sup>ad</sup> |
| Factory flock 1 | 6.35 +/- 0.14 <sup>d</sup> | 0.76 +/- 0.012 <sup>c</sup> | 16.4 +/- 0.7 <sup>b</sup> | 330 +/- 15 <sup>cd</sup> |
| Factory flock 2 | 5.80 +/- 0.18 <sup>b</sup> | 0.72 +/- 0.014 <sup>b</sup> | 16 +/- 0.9 <sup>a</sup> | 272 +/- 22 <sup>b</sup> |

To better understand the potential relationship between flock provenance and carriage of potential pathogens, we next focused on the relative abundance of taxa known to be of veterinary and medical importance by identifying ASVs that best matched known bird pathogens. V4 sequences representing *Staphylococcus sp*. were most prevalent in samples from commercially raised birds. *Staphylococcus* DNA was also detected in one flock of domestic free-ranging turkeys. Detectable levels of *Staphylococcus* DNA were not found in any samples from wild birds. Similarly, *Campylobacter* DNA was identified only in factory-farm raised birds. The abundance of *Campylobacter* DNA in some birds was suggestive of heavy colonization; however, it was undetected in other birds with in the same flock (Figure 2B, C,).

**Figure 2B, C, D.**
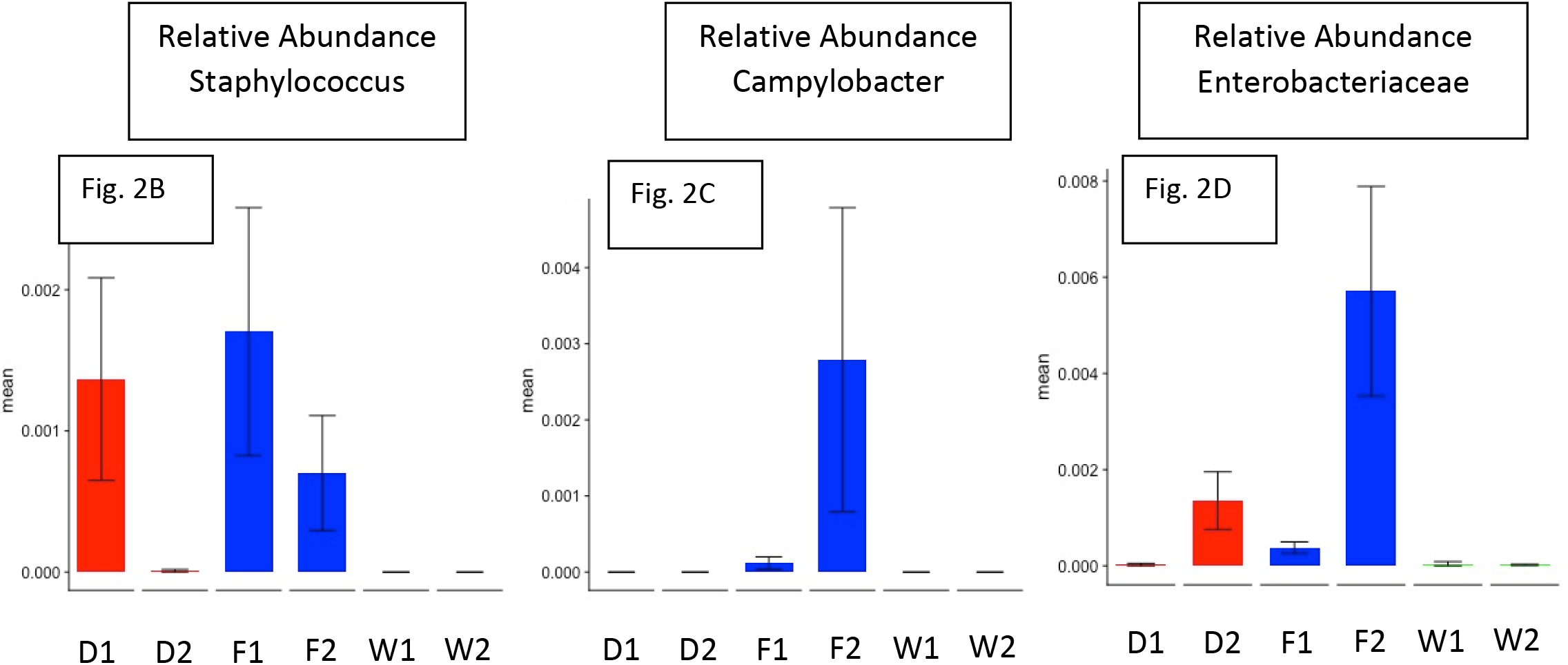
*Staphylococcus*, *Campylobacter* and Enterobacteriaceae are more abundant in the cecal contents of factory raised domestic turkeys than wild turkeys. Relative abundance of groups of ASVs (B, C) or an individual ASV (D) from the 16S sequencing dataset. Samples are grouped according to flock and colored by provenance: Red (D) = domestic birds raised on hobby farms; Blue (F) = Domestic birds raised in factory farms; Green (W) = free ranging wild turkeys)

One limitation of our approach is that without whole-genome data, the short region we sequenced cannot distinguish known pathogens from similar bacteria with identical sequences across the 16S V4 region. Measurable levels of the family *Enterobacteriaceae* were abundant in samples from both factory-raised flocks and one free-ranging domestic flock (Figure 2D). The *Enterobacteriaceae* are a large family of bacteria that include *Escherichia coli* as well as other pathogens including *Salmonell*a. The V4 region of the 16S rRNA gene does not resolve *E. coli* or *Salmonella* from other Enterobacteriaceae, which prevented us from estimating *E. coli* or *Salmonella* abundance in these animals through V4 sequencing alone. As *E. coli* and *Salmonell*a are common pathogens in domestic poultry production, we further investigated the prevalence of these potential pathogens in wild and factory-raised turkeys.

The presence of *Salmonella* DNA was detected by PCR targeting the *Salmonella* specific gene *invA* in total DNA isolated from cecal samples of individual birds. Of 14 samples tested from factory-raised birds, 11 tested positive for *invA*. Conversely, none of 11 samples collected from wild birds tested positive for the presence of the *invA* gene, suggesting that factory-raised turkeys more frequently contain *Salmonella* in their digestive tracts than wild turkeys. To determine the relative abundance of *E. coli* in cecal samples, a qPCR assay targeting the *E. coli* specific gene *ybbW* (42) was used. Genomic *E. coli* DNA was clearly present in the total DNA samples obtained from commercially-raised turkeys. Conversely, *E. coli* DNA in samples from wild turkeys was below the limit of detection of the assay (Figure 3). We also plated cecal samples on MacConkey agar to enrich for growth of enteric bacteria. Although not detectable by qPCR, we were able to isolate colonies characteristic of *E. coli* from wild turkey cecal samples. Their identity as *E. coli* was subsequently verified by amplification of the *ybbW* gene. *E. coli* were readily cultured from the ceca of factory-raised turkeys. In addition to colony growth consistent with *E. coli* (pink colonies), white colonies were also observed growing on MacConkey agar. These white colonies were not studied further or collected; however, based on growth on MacConkey agar, these colonies were likely *Salmonella* or other enteric bacteria.

**Figure 3.**
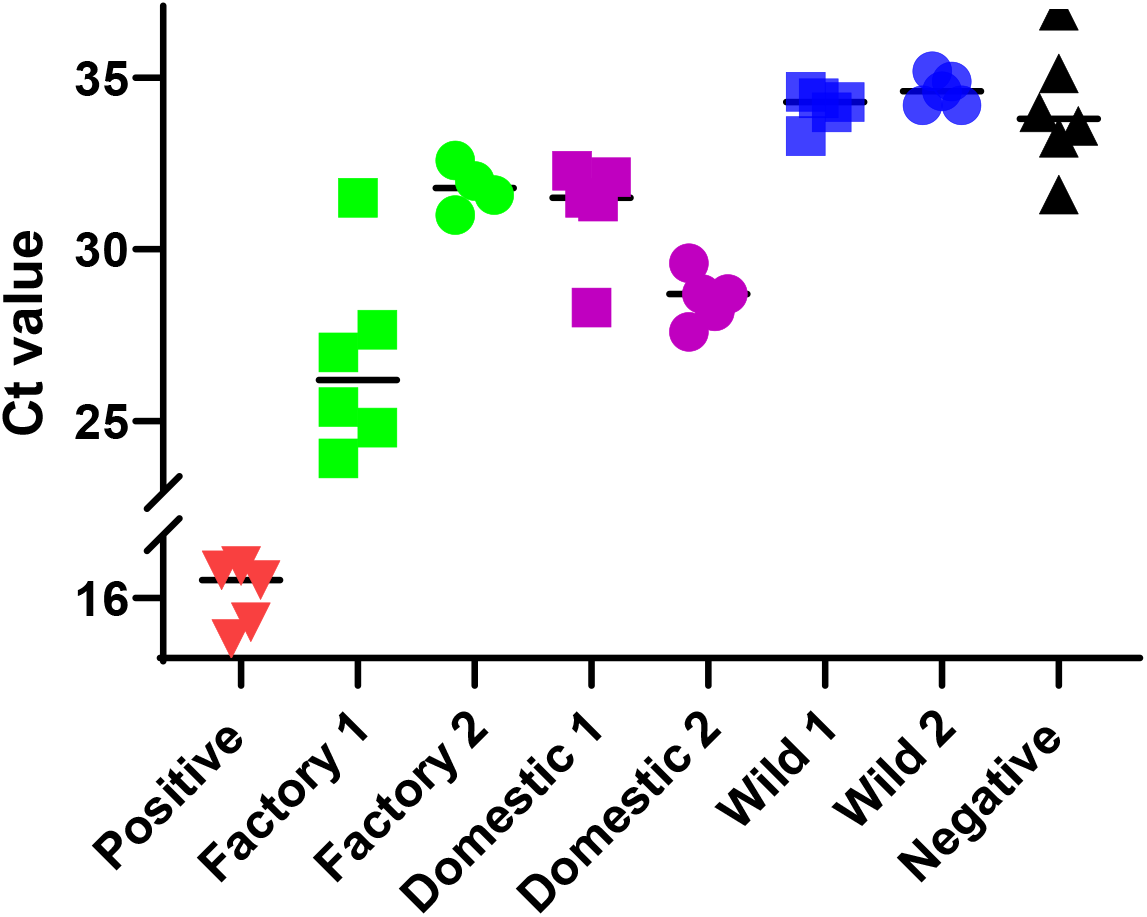
Cecal contents of factory raised and free ranging domestic turkeys contain high levels of *E. coli* DNA compared to wild birds. Quantitative PCR for the *ybbW* gene (*E. coli* specific) detected in total genomic DNA isolated from the ceca of factory-raised domestic, free-range domestic, and wild turkeys. CT values from individual birds are shown in the Y-axis. Average Ct values of each group are indicated by a horizontal bar. Individual flocks, positive control (100% *E. coli* genomic DNA) and negative control are indicated in the X-axis.

As we were able to isolate *E. coli* colonies from the ceca of both wild and factory-raised turkeys, we were interested in further understanding the differences that may exist between these bacterial populations. We therefor performed phylo-group analysis to compare the diversity of *E. coli* lineages that were isolated from factory-raised and wild turkeys. Of *E. coli* isolated from wild turkeys, 29 of 30 strains belonged to groups A, B1, or E whereas none belonged to groups B2, C, or D (Figure 4 and Supplementary Data S5). Strains isolated from factory-raised turkeys were more diverse with all major phylo-groups represented. Several (9/50) belonged to cryptic clades I or II, which have been infrequently isolated in other studies. Seven strains were classified as group B2 or D, which are lineages that are commonly associated with extraintestinal pathogenic *E. coli* strains (45, 46). These results suggested that the pathogenic potential of the *E. coli* present in wild turkeys may be different from those present in factory-raised domestic turkeys.

**Figure 4.**
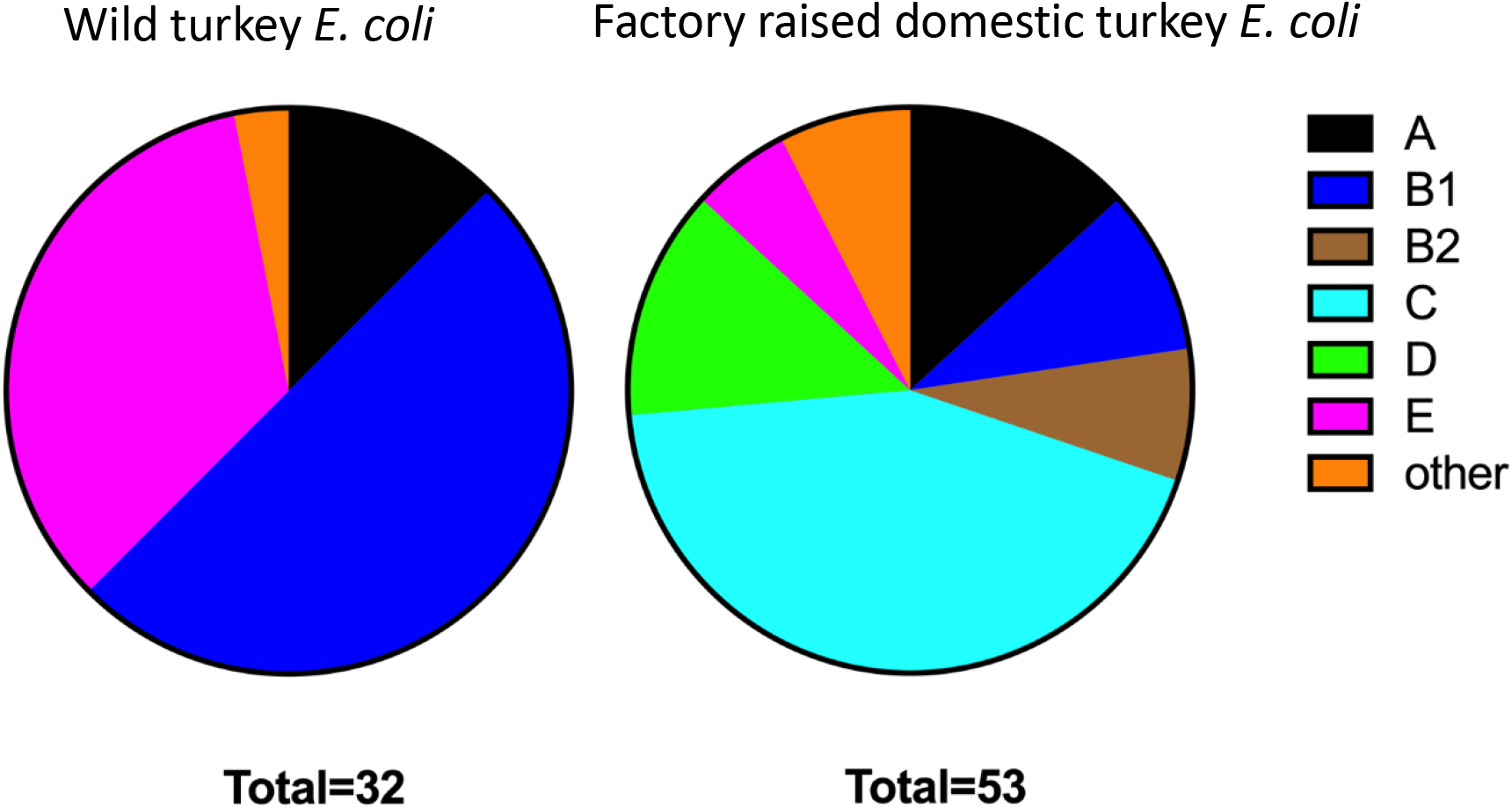
*E. coli* strains isolated from wild turkeys are predominantly phylo-groups B1 and E. A wide variety of phylo-groups populate the intestinal tract of factory raised turkeys Phylo-groups represented by color as indicated.

A number of virulence factors have been identified in extraintestinal pathogenic *E. coli*. These include proteins essential for iron acquisition and for group 2 or group 3 capsule production (47–53). To determine if virulence associated gene carriage differed between *E. coli* found in wild turkeys and factory-raised turkeys, end point PCR was used to determine carriage of three siderophore receptor genes (*iutA, iroN,* and *fyuA*) as well as the *kpsMT* genes involved in group 2 or group 3 capsule synthesis (Table 3). Nearly half (47%) of *E. coli* strains isolated from wild turkeys carried the aerobactin receptor *iutA* gene. However, carriage of the salmochelin receptor *iroN*, yersiniabactin receptor *fyuA*, or capsule synthesis *kpsMT* genes was not observed in *E. coli* strains isolated from wild turkeys. Conversely, only 10% of strains isolated from the ceca of commercially produced turkeys contained *iutA*, whereas *iroN* was present in 22% and *fyuA* in 4%. Capsule synthesis genes (*kpsMTII* or *kpsMTIII*) were present in 40% of strains isolated from factory-raised turkeys. Presence of virulence factors was not associated with any particular phylo-group, and several strains carried combinations of virulence factor genes (Supplemental Data S5).

**Table 3.**
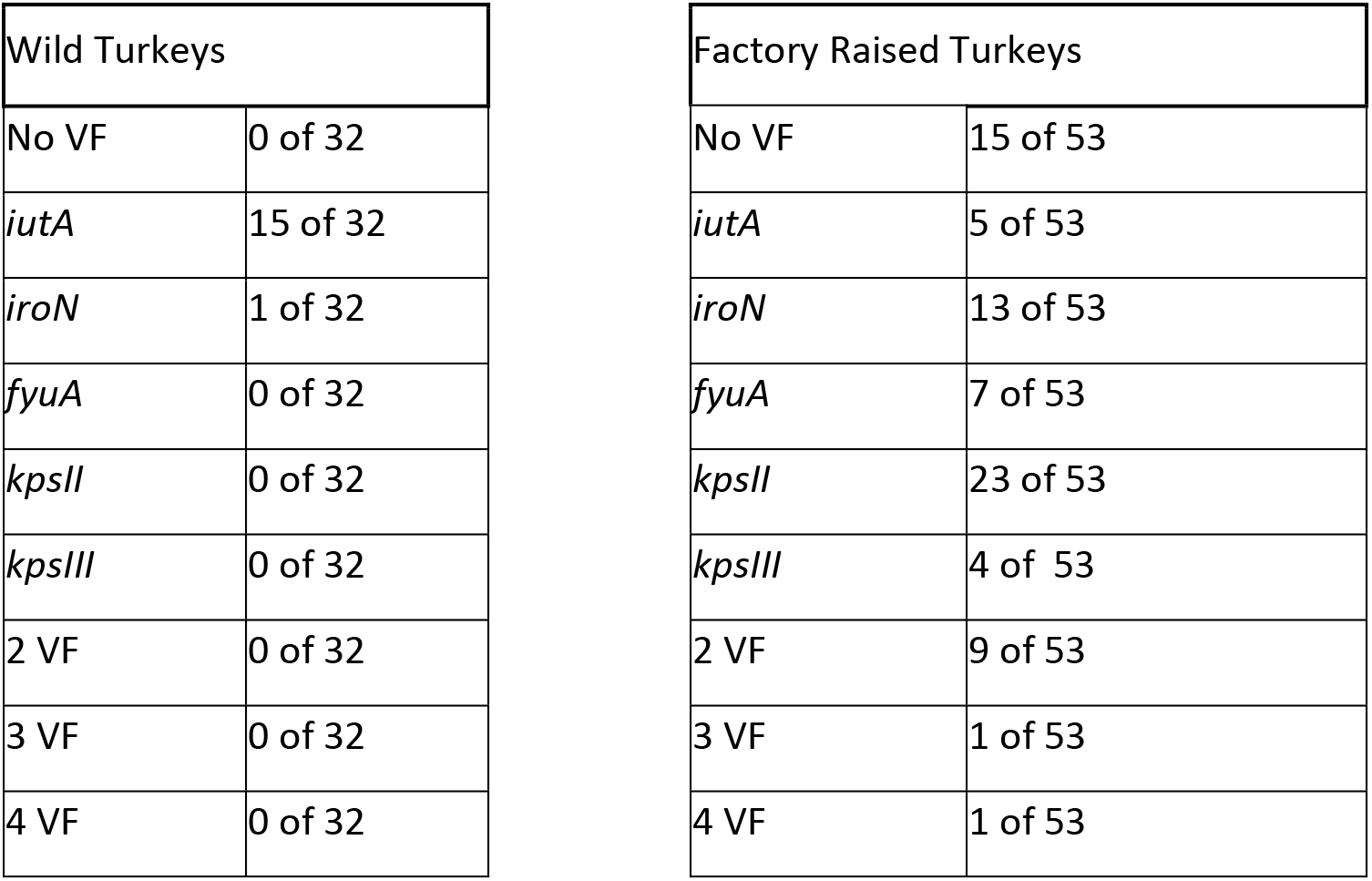
Prevalence of virulence factors genes in *E. coli* strains isolated from wild and factory raised turkeys. VF= virulence genes

## Discussion

The essential role of gut microbiota in maintaining animal and human health has been well established (54–57). Although diet is clearly an important selector for many functional guilds of microbes within the gut, host evolutionary history is thought to be a driving factor in determining the prevalence of specific microbial OTUs (58). Increasingly, evidence supports the theory that many animals coevolved with their microbial symbionts giving both host and microbe survival advantages (59–62). In addition to diet, the intestinal microbiome of domestic farm animals (including poultry) is likely influenced by a number of factors, such as, past and present exposure to antibiotics, exposure to the microbiome of the mother and other microbes in their environment. Data presented here suggest that common production practices (potentially in combination with) selective breeding in modern poultry farming have resulted in a turkey microbiome in which beta diversity decreases from wild birds to free-ranging domestic birds to the highly monotaxic microbiota seen in commercially raised turkeys. V4 sequencing results from this study are largely consistent with a previous clone-based sequencing approach comparing the microbiota of wild turkeys and domestic turkeys (11).

The composition of the gut microbiota in poultry likely influences a variety of beneficial characteristics, including immune system development and function (63–65). Domestic turkeys, raised in commercial turkey production facilities are highly susceptible to a myriad of economically devastating bacterial, fungal, viral and parasitic diseases (50). Previous research has shown that colonization by some commensal species of microbes prevents/inhibits colonization by pathogenic *Campylobacter, Staphylococcus* and *Salmonella* (66–73) in poultry. We hypothesize that wild relatives of agriculturally important species may carry a heritable microbiome which inhibits colonization by common pathogens. Modern agricultural production practices have largely ignored the potential benefits of this natural microbiota, having instead relied on widespread use of antibiotics to control pathogens.

While we have not yet established any specific mechanistic links between members of the normal flora and the abundance of specific pathogenic species, we observed that wild turkeys have higher levels of Coriobacteriales compared to hobby farm or factory-raised domestic turkeys. Some Coriobacteriales produce hydroxysteroid dehydrogenase enzymes involved in the conversion of primary to secondary bile acids (74, 75). Bile salt conversion has demonstrated effects on composition of the microbiome, colonization of intestinal pathogens, and immune responses in humans and livestock (74, 76–79). Growth of Coriobacteriales is stimulated by polyphenols found in diverse plants, and these bacteria metabolize them to phenolic compounds that have anti-inflammatory and immunomodulatory effects (75). Coriobacteriales are also especially prone to disruption by antibiotic treatment in mice (80). The diets of wild turkeys are free from agricultural antibiotics and likely contain diverse plant polyphenols. Whether members of this family are involved in colonization resistance to *Campylobacter*, *Salmonella*, or *E. coli* should be investigated further.

Suppression of avian-pathogenic *E. coli* in turkeys is an especially important priority for poultry producers. Therefore, it is notable that wild turkeys appeared to contain very little *E. coli* in their ceca. The *E. coli* strains isolated from wild turkeys were dissimilar to those isolated from factory-raised birds, both in terms of phylogenetic lineage as well as the presence of specific virulence factors. Many of the *E. coli* strains isolated from wild turkeys contained the aerobactin receptor gene. Aerobactin is a proven virulence factor in extraintestinal avian infections (51); however its role in these strains may be related to fitness in the highly competitive environment of the wild turkey intestinal tract. The absence of capsule synthesis genes, salmochelin, and yersiniabactin production from strains isolated from wild turkeys may indicate that these strains are not prone to cause bloodstream infections or colonize other organs. It is possible that bacteriocins, prophages, contact-dependent inhibition or type 6 secretion systems of *E. coli* lineages established wild turkeys exclude invasion by avian-pathogenic strains frequently found in factory-raised poultry (81–84).

Due to common poultry production practices, microbes colonizing the intestinal tract of commercially raised poultry are minimally, if at all, influenced by the microbiome of the mother. The practice of hatching surface sterilized eggs in incubators for multiple generations has surely contributed to the loss of heritable microbial taxa which coevolved with the wild turkey over millennia. As a consequence, modern production practices have likely resulted in domestic poultry obtaining their microbiota almost exclusively from the environment found in the production facilities in which they are raised. These conditions have likely skewed the gut microbiota toward taxa best capable of survival in modern turkey production facilities rather than taxa contributing to the mutual survival of host and microbe.

The transfer of microbiota from mother to infant has been best characterized in mammals. The transfer of maternal microbes to the mammalian young begins during the birthing process and continues through nursing and social interactions (85, 86). Coprophagy is common in many animal species, including turkeys (87–89) and it is common to see turkeys consuming cecal drops of their cage mates. This innate behavior in turkeys may have evolved to enable bird-to-bird spread of beneficial microbiota within a flock. Recent work documents that the newly hatched young of some birds readily consume cecal drops, but not normal rectal feces, of their mothers. This consumption of maternal cecal drops by chicks was observed only during a short window of time (approximately the first month of life) (90). This behavior potentially facilitates the establishment of a beneficial, heritable gut microbiome from mother to chick.

The gut microbiome is perhaps one of the most complex of biological communities. As in the analysis of any biological community, it is essential to consider the effects of dominant taxa, as well as taxa that may comprise a relatively small, but potentially important, role in the community as a whole. The goal of this study was to identify potential changes/differences in the microbial composition of factory-raised turkeys when compared to their wild predecessors. Results presented here demonstrate that the overall abundance of *E. coli*, *Salmonella, Campylobacter* and *Staphylococcus* in wild turkeys is much lower than levels commonly found in commercially raised turkeys. Furthermore, *E. coli* strains occupying the intestinal tract of wild turkeys appear distinct from strains commonly found in commercially raised turkeys. The strong correlation between bird provenance, increased microbial diversity and low pathogen carriage warrants further research into the potential for mining the microbiome of free-ranging wild turkeys (as well as wild relatives of other agriculturally important species) in search of therapeutics or probiotics for use in controlling pathogens common in agricultural food production.

## Supplemental Data Figure S1

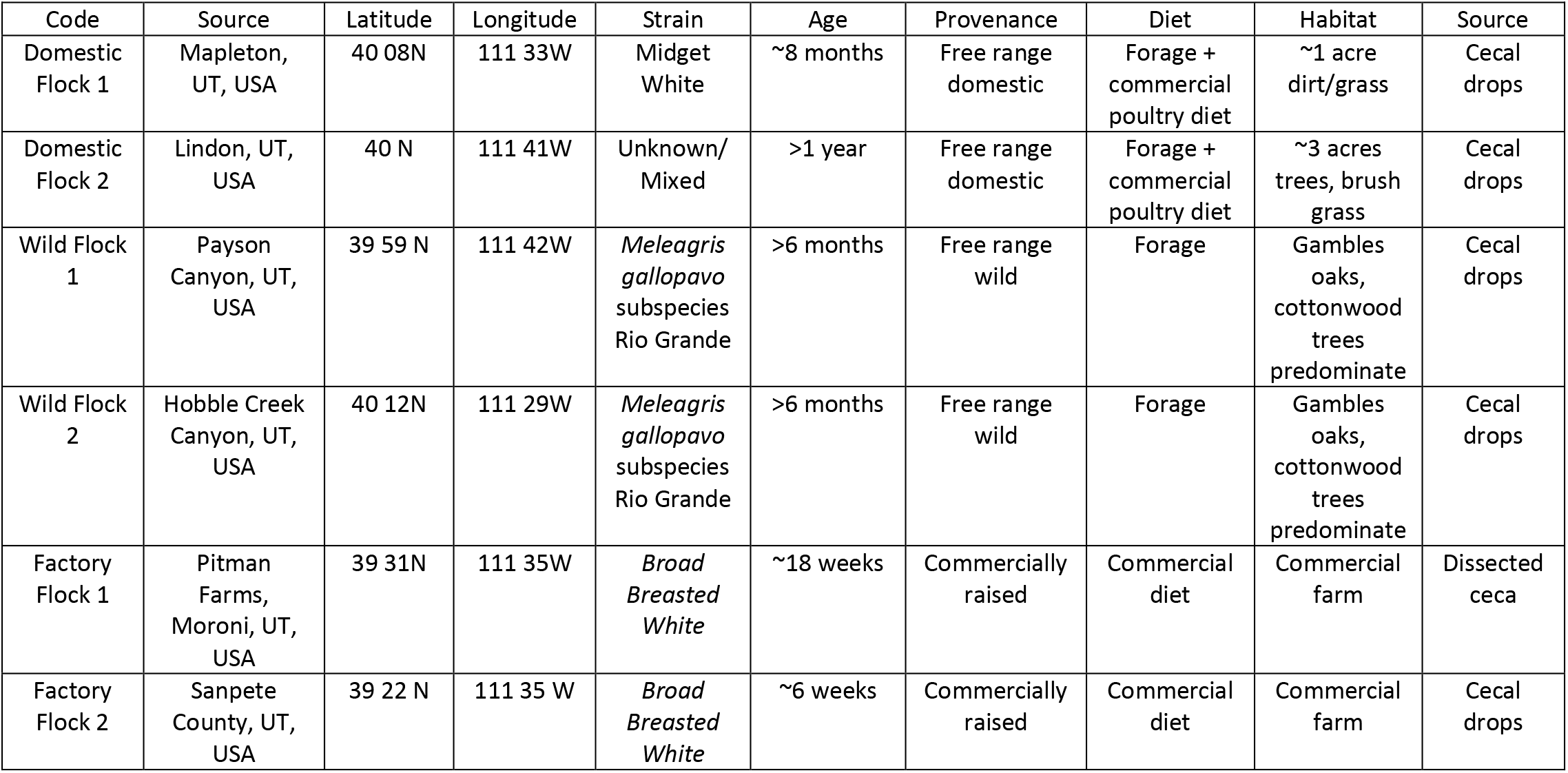

## Supplemental Data S2

### qPCR

For qPCR experiments, total DNA was isolated from cecal samples using the Qiagen blood and tissue DNA kit as directed by the manufacturer. DNA samples were diluted to a concentration of 100ng/μl total DNA. IDT PrimeTime^®^ Gene Expression Master Mix was used in all qPCR assays. PCR conditions: Briefly, thermocycling was performed as suggested by manufacture (40 cycles of 95 °C denaturation for 15 seconds followed 57 °C annealing/amplification for one minute) using an ABI StepOnePlus™ Real-Time PCR System. qPCR primers and probe were designed using IDT PrimerQuest^®^ and manufactured by Integrated DNA Technologies.

<u>Primer and probe sequences for *ybbW*</u>

Primer 1F TGATTGGCAAATCTGGCCG

Primer 1R CGTTGACCAGCCAGAAGATTAAG

Probe 56-FAM/AAGCCCGGT/ZEN/AGAGAAAGGCCTAAC/3IABkFQ\

### *invA* Detection

In endpoint PCR experiments to detect the presence of the *invA* gene, OneTaq master mix (NEB) was used in all reactions. The following PCR conditions were used: 95 °C for 2 minutes followed by 30 cycles of 95 °C for 20 sec. 56 °C for 20 sec. 68 °C for 40 sec. followed by a 7 minute final extension step at 68 °C. Amplified DNA was visualized by running the product on a 1.75% agarose gel.

<u>>Primer sequences for *invA*</u>

*invA* Primer 1F GTGAAATTATCGCCACGTTCGGGCAA

*invA* Primer 1R TCATCGCACCGTCAAAGGAACC

### Phylo-Group Determination

Phylogrouping was performed as previously described (1) except that we used 10 μl reactions in OneTaq master mix (NEB) with an extension temperature of 68 degrees. We first performed the quadriplex PCR reaction, followed by group E or group C specific PCR when warranted to distinguish between D/E or A/C strains.

Virulence genotyping was performed using approximately 100 ng genomic DNA as template and 20 pmol of each primer in OneTaq master mix. The conditions were 94 °C for 3 min followed by 30 cycles of 94 °C for 15 seconds, 57 °C for 15 seconds, and 68 °C for 45 seconds, and a final extension of 68 °C for 5 min. The *fyuA, kpsII* and *kpsIII* reactions were multiplexed, the *iutA iroN* and *iutA* reactions were run individually.

Primer Names, Sequences, Target and Expected Amplicon Size.

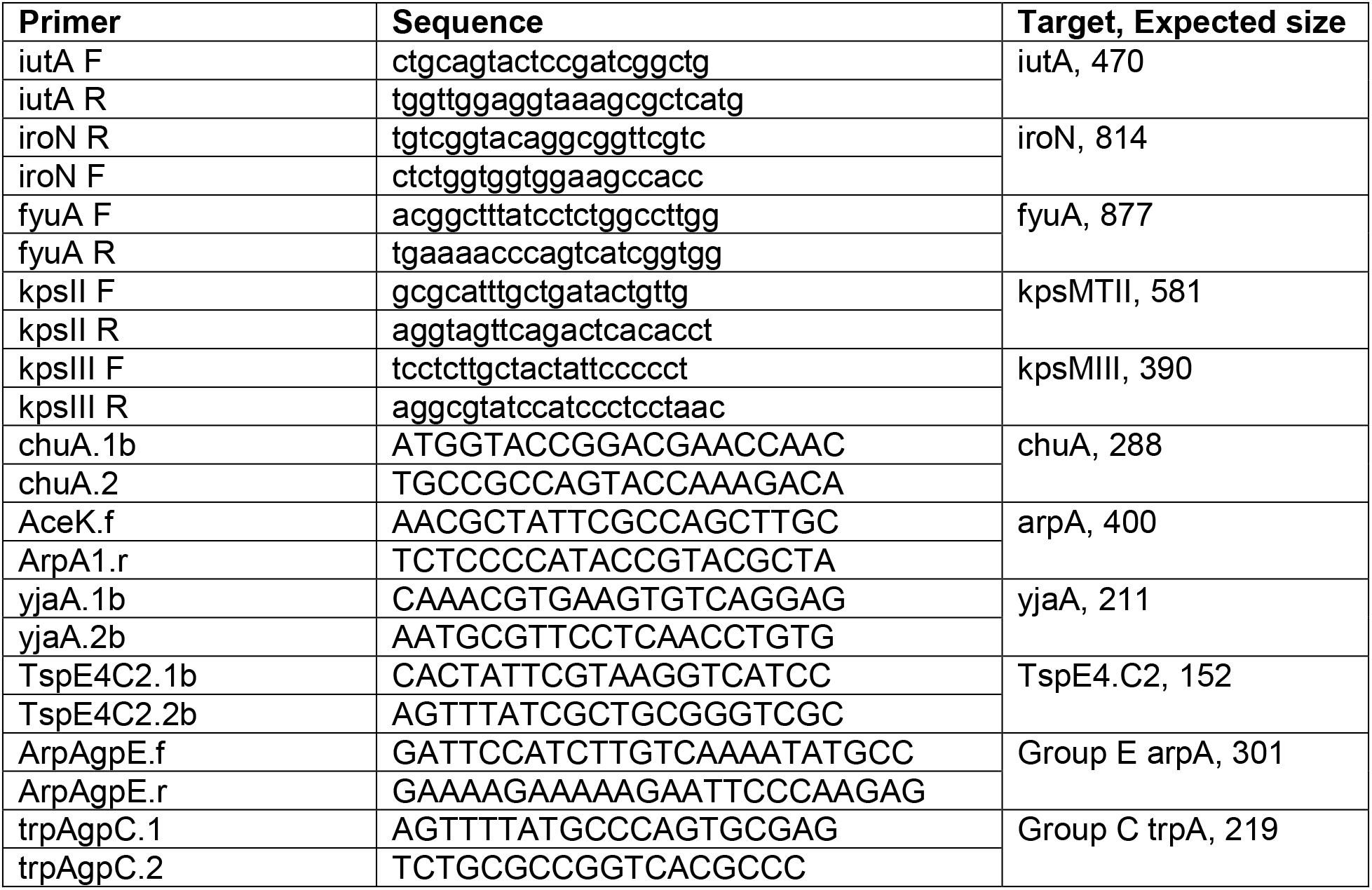

## References

1. Speller CF, Kemp BM, Wyatt SD, Monroe C, Lipe WD, Arndt UM, Yang DY. 2010. Ancient mitochondrial DNA analysis reveals complexity of indigenous North American turkey domestication. Proc Natl Acad Sci U S A 107:2807–12.

2. Guan X, Silva P, Gyenai KB, Xu J, Geng T, Tu Z, Samuels DC, Smith EJ. 2009. The mitochondrial genome sequence and molecular phylogeny of the turkey, Meleagris gallopavo. Anim Genet 40:134–41.

3. Thornton EK, Emery KF, Steadman DW, Speller C, Matheny R, Yang D. 2012. Earliest Mexican Turkeys (Meleagris gallopavo) in the Maya Region: implications for pre-Hispanic animal trade and the timing of turkey domestication. PLoS One 7:e42630.

4. Aslam ML, Bastiaansen JW, Elferink MG, Megens HJ, Crooijmans RP, Blomberg le A, Fleischer RC, Van Tassell CP, Sonstegard TS, Schroeder SG, Groenen MA, Long JA. 2012. Whole genome SNP discovery and analysis of genetic diversity in Turkey (Meleagris gallopavo). BMC Genomics 13:391.

5. Canales Vergara AM, Landi V, Delgado Bermejo JV, Martinez A, Cervantes Acosta P, Pons Barro A, Bigi D, Sponenberg P, Helal M, Hossein Banabazi M, Camacho Vallejo ME. 2019. Tracing Worldwide Turkey Genetic Diversity Using D-loop Sequence Mitochondrial DNA Analysis. Animals (Basel) 9.

6. Stover KK, Weinreich DM, Roberts TJ, Brainerd EL. 2018. Patterns of musculoskeletal growth and dimensional changes associated with selection and developmental plasticity in domestic and wild strain turkeys. Ecol Evol 8:3229–3239.

7. USDA. Turkey Sector: Background & Statistics.

8. Colston TJ, Jackson CR. 2016. Microbiome evolution along divergent branches of the vertebrate tree of life: what is known and unknown. Mol Ecol 25:3776–800.

9. Clayton JB, Vangay P, Huang H, Ward T, Hillmann BM, Al-Ghalith GA, Travis DA, Long HT, Tuan BV, Minh VV, Cabana F, Nadler T, Toddes B, Murphy T, Glander KE, Johnson TJ, Knights D. 2016. Captivity humanizes the primate microbiome. Proc Natl Acad Sci U S A 113:10376–81.

10. McKenzie VJ, Song SJ, Delsuc F, Prest TL, Oliverio AM, Korpita TM, Alexiev A, Amato KR, Metcalf JL, Kowalewski M, Avenant NL, Link A, Di Fiore A, Seguin-Orlando A, Feh C, Orlando L, Mendelson JR, Sanders J, Knight R. 2017. The Effects of Captivity on the Mammalian Gut Microbiome. Integr Comp Biol 57:690–704.

11. Scupham AJ, Patton TG, Bent E, Bayles DO. 2008. Comparison of the cecal microbiota of domestic and wild turkeys. Microb Ecol 56:322–31.

12. Schreuder J, Velkers FC, Bouwstra RJ, Beerens N, Stegeman JA, de Boer WF, van Hooft P, Elbers ARW, Bossers A, Jurburg SD. 2020. An observational field study of the cloacal microbiota in adult laying hens with and without access to an outdoor range. Anim Microbiome 2:28.

13. Cui Y, Wang Q, Liu S, Sun R, Zhou Y, Li Y. 2017. Age-Related Variations in Intestinal Microflora of Free-Range and Caged Hens. Front Microbiol 8:1310.

14. Hubert SM, Al-Ajeeli M, Bailey CA, Athrey G. 2019. The Role of Housing Environment and Dietary Protein Source on the Gut Microbiota of Chicken. Animals (Basel) 9.

15. Souza MR, Moreira JL, Barbosa FH, Cerqueira MM, Nunes AC, Nicoli JR. 2007. Influence of intensive and extensive breeding on lactic acid bacteria isolated from Gallus gallus domesticus ceca. Vet Microbiol 120:142–50.

16. Pascoe EL, Hauffe HC, Marchesi JR, Perkins SE. 2017. Network analysis of gut microbiota literature: an overview of the research landscape in non-human animal studies. ISME J 11:2644–2651.

17. Hird SM. 2017. Evolutionary Biology Needs Wild Microbiomes. Front Microbiol 8:725.

18. Brisbin JT, Gong J, Sharif S. 2008. Interactions between commensal bacteria and the gut-associated immune system of the chicken. Anim Health Res Rev 9:101–10.

19. Yeoman CJ, Chia N, Jeraldo P, Sipos M, Goldenfeld ND, White BA. 2012. The microbiome of the chicken gastrointestinal tract. Anim Health Res Rev 13:89–99.

20. Danzeisen JL, Calvert AJ, Noll SL, McComb B, Sherwood JS, Logue CM, Johnson TJ. 2013. Succession of the turkey gastrointestinal bacterial microbiome related to weight gain. PeerJ 1:e237.

21. Wei S, Gutek A, Lilburn M, Yu Z. 2013. Abundance of pathogens in the gut and litter of broiler chickens as affected by bacitracin and litter management. Vet Microbiol 166:595–601.

22. Taylor KJM, Ngunjiri JM, Abundo MC, Jang H, Elaish M, Ghorbani A, Kc M, Weber BP, Johnson TJ, Lee CW. 2020. Respiratory and Gut Microbiota in Commercial Turkey Flocks with Disparate Weight Gain Trajectories Display Differential Compositional Dynamics. Appl Environ Microbiol 86.

23. Danzeisen JL, Clayton JB, Huang H, Knights D, McComb B, Hayer SS, Johnson TJ. 2015. Temporal Relationships Exist Between Cecum, Ileum, and Litter Bacterial Microbiomes in a Commercial Turkey Flock, and Subtherapeutic Penicillin Treatment Impacts Ileum Bacterial Community Establishment. Front Vet Sci 2:56.

24. Johnson TA, Sylte MJ, Looft T. 2019. In-feed bacitracin methylene disalicylate modulates the turkey microbiota and metabolome in a dose-dependent manner. Sci Rep 9:8212.

25. Wilkinson TJ, Cowan AA, Vallin HE, Onime LA, Oyama LB, Cameron SJ, Gonot C, Moorby JM, Waddams K, Theobald VJ, Leemans D, Bowra S, Nixey C, Huws SA. 2017. Characterization of the Microbiome along the Gastrointestinal Tract of Growing Turkeys. Front Microbiol 8:1089.

26. Fenna L, Boag DA. 1974. Filling and emptying of the galliform caecum. Can J Zool 52:537–40.

27. Barnes EM, Mead GC, Barnum DA, Harry EG. 1972. The intestinal flora of the chicken in the period 2 to 6 weeks of age, with particular reference to the anaerobic bacteria. Br Poult Sci 13:311–26.

28. Bjerrum L, Engberg RM, Leser TD, Jensen BB, Finster K, Pedersen K. 2006. Microbial community composition of the ileum and cecum of broiler chickens as revealed by molecular and culture-based techniques. Poult Sci 85:1151–64.

29. Pauwels J, Taminiau B, Janssens GP, De Beenhouwer M, Delhalle L, Daube G, Coopman F. 2015. Cecal drop reflects the chickens’ cecal microbiome, fecal drop does not. J Microbiol Methods 117:164–70.

30. Kozich JJ, Westcott SL, Baxter NT, Highlander SK, Schloss PD. 2013. Development of a dual-index sequencing strategy and curation pipeline for analyzing amplicon sequence data on the MiSeq Illumina sequencing platform. Appl Environ Microbiol 79:5112–20.

31. Bolyen E, Rideout JR, Dillon MR, Bokulich NA, Abnet CC, Al-Ghalith GA, Alexander H, Alm EJ, Arumugam M, Asnicar F, Bai Y, Bisanz JE, Bittinger K, Brejnrod A, Brislawn CJ, Brown CT, Callahan BJ, Caraballo-Rodriguez AM, Chase J, Cope EK, Da Silva R, Diener C, Dorrestein PC, Douglas GM, Durall DM, Duvallet C, Edwardson CF, Ernst M, Estaki M, Fouquier J, Gauglitz JM, Gibbons SM, Gibson DL, Gonzalez A, Gorlick K, Guo J, Hillmann B, Holmes S, Holste H, Huttenhower C, Huttley GA, Janssen S, Jarmusch AK, Jiang L, Kaehler BD, Kang KB, Keefe CR, Keim P, Kelley ST, Knights D, et al. 2019. Author Correction: Reproducible, interactive, scalable and extensible microbiome data science using QIIME 2. Nat Biotechnol 37:1091.

32. Bolyen E, Rideout JR, Dillon MR, Bokulich NA, Abnet CC, Al-Ghalith GA, Alexander H, Alm EJ, Arumugam M, Asnicar F, Bai Y, Bisanz JE, Bittinger K, Brejnrod A, Brislawn CJ, Brown CT, Callahan BJ, Caraballo-Rodriguez AM, Chase J, Cope EK, Da Silva R, Diener C, Dorrestein PC, Douglas GM, Durall DM, Duvallet C, Edwardson CF, Ernst M, Estaki M, Fouquier J, Gauglitz JM, Gibbons SM, Gibson DL, Gonzalez A, Gorlick K, Guo J, Hillmann B, Holmes S, Holste H, Huttenhower C, Huttley GA, Janssen S, Jarmusch AK, Jiang L, Kaehler BD, Kang KB, Keefe CR, Keim P, Kelley ST, Knights D, et al. 2019. Reproducible, interactive, scalable and extensible microbiome data science using QIIME 2. Nat Biotechnol 37:852–857.

33. Callahan BJ, McMurdie PJ, Rosen MJ, Han AW, Johnson AJ, Holmes SP. 2016. DADA2: High-resolution sample inference from Illumina amplicon data. Nat Methods 13:581–3.

34. McDonald D, Price MN, Goodrich J, Nawrocki EP, DeSantis TZ, Probst A, Andersen GL, Knight R, Hugenholtz P. 2012. An improved Greengenes taxonomy with explicit ranks for ecological and evolutionary analyses of bacteria and archaea. ISME J 6:610–8.

35. Oksanen J, Blanchet, F. G., Friendly, M., Kindt, R., Legendre, P., McGlinn, D., . . . Wagner, H. . 2018. vegan: Community Ecology Package.

36. Lozupone CA, Hamady M, Kelley ST, Knight R. 2007. Quantitative and qualitative beta diversity measures lead to different insights into factors that structure microbial communities. Appl Environ Microbiol 73:1576–85.

37. Lozupone C, Knight R. 2005. UniFrac: a new phylogenetic method for comparing microbial communities. Appl Environ Microbiol 71:8228–35.

38. Price MN, Dehal PS, Arkin AP. 2010. FastTree 2--approximately maximum-likelihood trees for large alignments. PLoS One 5:e9490.

39. Katoh K, Misawa K, Kuma K, Miyata T. 2002. MAFFT: a novel method for rapid multiple sequence alignment based on fast Fourier transform. Nucleic Acids Res 30:3059–66.

40. Mandal S, Van Treuren W, White RA, Eggesbo M, Knight R, Peddada SD. 2015. Analysis of composition of microbiomes: a novel method for studying microbial composition. Microb Ecol Health Dis 26:27663.

41. Bokulich NA, Kaehler BD, Rideout JR, Dillon M, Bolyen E, Knight R, Huttley GA, Gregory Caporaso J. 2018. Optimizing taxonomic classification of marker-gene amplicon sequences with QIIME 2’s q2-feature-classifier plugin. Microbiome 6:90.

42. Walker DI, McQuillan J, Taiwo M, Parks R, Stenton CA, Morgan H, Mowlem MC, Lees DN. 2017. A highly specific Escherichia coli qPCR and its comparison with existing methods for environmental waters. Water Res 126:101–110.

43. Clermont O, Christenson JK, Denamur E, Gordon DM. 2013. The Clermont Escherichia coli phylo-typing method revisited: improvement of specificity and detection of new phylo-groups. Environ Microbiol Rep 5:58–65.

44. Rahn K, De Grandis SA, Clarke RC, McEwen SA, Galan JE, Ginocchio C, Curtiss R, 3rd, Gyles CL. 1992. Amplification of an invA gene sequence of Salmonella typhimurium by polymerase chain reaction as a specific method of detection of Salmonella. Mol Cell Probes 6:271–9.

45. Picard B, Garcia JS, Gouriou S, Duriez P, Brahimi N, Bingen E, Elion J, Denamur E. 1999. The link between phylogeny and virulence in Escherichia coli extraintestinal infection. Infect Immun 67:546–53.

46. Johnson JR, Stell AL. 2000. Extended virulence genotypes of Escherichia coli strains from patients with urosepsis in relation to phylogeny and host compromise. J Infect Dis 181:261–72.

47. Dziva F, Stevens MP. 2008. Colibacillosis in poultry: unravelling the molecular basis of virulence of avian pathogenic Escherichia coli in their natural hosts. Avian Pathol 37:355–66.

48. Mageiros L, Meric G, Bayliss SC, Pensar J, Pascoe B, Mourkas E, Calland JK, Yahara K, Murray S, Wilkinson TS, Williams LK, Hitchings MD, Porter J, Kemmett K, Feil EJ, Jolley KA, Williams NJ, Corander J, Sheppard SK. 2021. Genome evolution and the emergence of pathogenicity in avian Escherichia coli. Nat Commun 12:765.

49. La Ragione RM, Woodward MJ. 2002. Virulence factors of Escherichia coli serotypes associated with avian colisepticaemia. Res Vet Sci 73:27–35.

50. Swayne DE. 2020. Diseases of poultry, Fourteenth edition. ed. Wiley-Blackwell, Hoboken, NJ.

51. Dozois CM, Daigle F, Curtiss R, 3rd. 2003. Identification of pathogen-specific and conserved genes expressed in vivo by an avian pathogenic Escherichia coli strain. Proc Natl Acad Sci U S A 100:247–52.

52. Collingwood C, Kemmett K, Williams N, Wigley P. 2014. Is the Concept of Avian Pathogenic Escherichia coli as a Single Pathotype Fundamentally Flawed? Front Vet Sci 1:5.

53. Antao EM, Glodde S, Li G, Sharifi R, Homeier T, Laturnus C, Diehl I, Bethe A, Philipp HC, Preisinger R, Wieler LH, Ewers C. 2008. The chicken as a natural model for extraintestinal infections caused by avian pathogenic Escherichia coli (APEC). Microb Pathog 45:361–9.

54. Clemente JC, Ursell LK, Parfrey LW, Knight R. 2012. The impact of the gut microbiota on human health: an integrative view. Cell 148:1258–70.

55. Sekirov I, Russell SL, Antunes LC, Finlay BB. 2010. Gut microbiota in health and disease. Physiol Rev 90:859–904.

56. Peixoto RS, Harkins DM, Nelson KE. 2021. Advances in Microbiome Research for Animal Health. Annu Rev Anim Biosci 9:289–311.

57. Waite DW, Taylor MW. 2014. Characterizing the avian gut microbiota: membership, driving influences, and potential function. Front Microbiol 5:223.

58. Youngblut ND, Reischer GH, Walters W, Schuster N, Walzer C, Stalder G, Ley RE, Farnleitner AH. 2019. Host diet and evolutionary history explain different aspects of gut microbiome diversity among vertebrate clades. Nat Commun 10:2200.

59. Moeller AH, Li Y, Mpoudi Ngole E, Ahuka-Mundeke S, Lonsdorf EV, Pusey AE, Peeters M, Hahn BH, Ochman H. 2014. Rapid changes in the gut microbiome during human evolution. Proc Natl Acad Sci U S A 111:16431–5.

60. Moeller AH, Caro-Quintero A, Mjungu D, Georgiev AV, Lonsdorf EV, Muller MN, Pusey AE, Peeters M, Hahn BH, Ochman H. 2016. Cospeciation of gut microbiota with hominids. Science 353:380–2.

61. Groussin M, Mazel F, Alm EJ. 2020. Co-evolution and Co-speciation of Host-Gut Bacteria Systems. Cell Host Microbe 28:12–22.

62. Ley RE, Hamady M, Lozupone C, Turnbaugh PJ, Ramey RR, Bircher JS, Schlegel ML, Tucker TA, Schrenzel MD, Knight R, Gordon JI. 2008. Evolution of mammals and their gut microbes. Science 320:1647–51.

63. Mwangi WN, Beal RK, Powers C, Wu X, Humphrey T, Watson M, Bailey M, Friedman A, Smith AL. 2010. Regional and global changes in TCRalphabeta T cell repertoires in the gut are dependent upon the complexity of the enteric microflora. Dev Comp Immunol 34:406–17.

64. van Veelen HPJ, Falcao Salles J, Matson KD, van der Velde M, Tieleman BI. 2020. Microbial environment shapes immune function and cloacal microbiota dynamics in zebra finches Taeniopygia guttata. Anim Microbiome 2:21.

65. McFall-Ngai M. 2007. Adaptive immunity: care for the community. Nature 445:153.

66. Cole K, Farnell MB, Donoghue AM, Stern NJ, Svetoch EA, Eruslanov BN, Volodina LI, Kovalev YN, Perelygin VV, Mitsevich EV, Mitsevich IP, Levchuk VP, Pokhilenko VD, Borzenkov VN, Svetoch OE, Kudryavtseva TY, Reyes-Herrera I, Blore PJ, Solis de los Santos F, Donoghue DJ. 2006. Bacteriocins reduce Campylobacter colonization and alter gut morphology in turkey poults. Poult Sci 85:1570–5.

67. Stern NJ, Svetoch EA, Eruslanov BV, Kovalev YN, Volodina LI, Perelygin VV, Mitsevich EV, Mitsevich IP, Levchuk VP. 2005. Paenibacillus polymyxa purified bacteriocin to control Campylobacter jejuni in chickens. J Food Prot 68:1450–3.

68. Stern NJ, Svetoch EA, Eruslanov BV, Perelygin VV, Mitsevich EV, Mitsevich IP, Pokhilenko VD, Levchuk VP, Svetoch OE, Seal BS. 2006. Isolation of a Lactobacillus salivarius strain and purification of its bacteriocin, which is inhibitory to Campylobacter jejuni in the chicken gastrointestinal system. Antimicrob Agents Chemother 50:3111–6.

69. Scupham AJ, Jones JA, Rettedal EA, Weber TE. 2010. Antibiotic manipulation of intestinal microbiota to identify microbes associated with Campylobacter jejuni exclusion in poultry. Appl Environ Microbiol 76:8026–32.

70. Scupham AJ. 2009. Campylobacter colonization of the Turkey intestine in the context of microbial community development. Appl Environ Microbiol 75:3564–71.

71. Nicoll TR, Jensen MM. 1987. Staphylococcosis of turkeys. 5. Large-scale control programs using bacterial interference. Avian Dis 31:85–8.

72. Wilkinson DM, Jensen MM. 1987. Staphylococcosis of turkeys. 4. Characterization of a bacteriocin produced by an interfering Staphylococcus. Avian Dis 31:80–4.

73. Bielke LR, Elwood AL, Donoghue DJ, Donoghue AM, Newberry LA, Neighbor NK, Hargis BM. 2003. Approach for selection of individual enteric bacteria for competitive exclusion in turkey poults. Poult Sci 82:1378–82.

74. Ridlon JM, Kang DJ, Hylemon PB. 2006. Bile salt biotransformations by human intestinal bacteria. J Lipid Res 47:241–59.

75. Rodriguez-Daza MC, Roquim M, Dudonne S, Pilon G, Levy E, Marette A, Roy D, Desjardins Y. 2020. Berry Polyphenols and Fibers Modulate Distinct Microbial Metabolic Functions and Gut Microbiota Enterotype-Like Clustering in Obese Mice. Front Microbiol 11:2032.

76. Gaboriaud P, Sadrin G, Guitton E, Fort G, Niepceron A, Lallier N, Rossignol C, Larcher T, Sausset A, Guabiraba R, Silvestre A, Lacroix-Lamande S, Schouler C, Laurent F, Bussiere FI. 2020. The Absence of Gut Microbiota Alters the Development of the Apicomplexan Parasite Eimeria tenella. Front Cell Infect Microbiol 10:632556.

77. McKenney PT, Yan J, Vaubourgeix J, Becattini S, Lampen N, Motzer A, Larson PJ, Dannaoui D, Fujisawa S, Xavier JB, Pamer EG. 2019. Intestinal Bile Acids Induce a Morphotype Switch in Vancomycin-Resistant Enterococcus that Facilitates Intestinal Colonization. Cell Host Microbe 25:695–705 e5.

78. Zhang L, Wu W, Lee YK, Xie J, Zhang H. 2018. Spatial Heterogeneity and Co-occurrence of Mucosal and Luminal Microbiome across Swine Intestinal Tract. Front Microbiol 9:48.

79. van Best N, Rolle-Kampczyk U, Schaap FG, Basic M, Olde Damink SWM, Bleich A, Savelkoul PHM, von Bergen M, Penders J, Hornef MW. 2020. Bile acids drive the newborn's gut microbiota maturation. Nat Commun 11:3692.

80. Zakostelska Z, Malkova J, Klimesova K, Rossmann P, Hornova M, Novosadova I, Stehlikova Z, Kostovcik M, Hudcovic T, Stepankova R, Juzlova K, Hercogova J, Tlaskalova-Hogenova H, Kverka M. 2016. Intestinal Microbiota Promotes Psoriasis-Like Skin Inflammation by Enhancing Th17 Response. PLoS One 11:e0159539.

81. Aoki SK, Pamma R, Hernday AD, Bickham JE, Braaten BA, Low DA. 2005. Contact-dependent inhibition of growth in Escherichia coli. Science 309:1245–8.

82. Journet L, Cascales E. 2016. The Type VI Secretion System in Escherichia coli and Related Species. EcoSal Plus 7.

83. Cascales E, Buchanan SK, Duche D, Kleanthous C, Lloubes R, Postle K, Riley M, Slatin S, Cavard D. 2007. Colicin biology. Microbiol Mol Biol Rev 71:158–229.

84. Da Re S, Valle J, Charbonnel N, Beloin C, Latour-Lambert P, Faure P, Turlin E, Le Bouguenec C, Renauld-Mongenie G, Forestier C, Ghigo JM. 2013. Identification of commensal Escherichia coli genes involved in biofilm resistance to pathogen colonization. PLoS One 8:e61628.

85. Dominguez-Bello MG, Costello EK, Contreras M, Magris M, Hidalgo G, Fierer N, Knight R. 2010. Delivery mode shapes the acquisition and structure of the initial microbiota across multiple body habitats in newborns. Proc Natl Acad Sci U S A 107:11971–5.

86. Mueller NT, Bakacs E, Combellick J, Grigoryan Z, Dominguez-Bello MG. 2015. The infant microbiome development: mom matters. Trends Mol Med 21:109–17.

87. Soave O, Brand CD. 1991. Coprophagy in animals: a review. Cornell Vet 81:357–64.

88. Pan D, Yu Z. 2014. Intestinal microbiome of poultry and its interaction with host and diet. Gut Microbes 5:108–19.

89. Kers JG, Velkers FC, Fischer EAJ, Hermes GDA, Stegeman JA, Smidt H. 2018. Host and Environmental Factors Affecting the Intestinal Microbiota in Chickens. Front Microbiol 9:235.

90. Kobayashi A, Tsuchida S, Ueda A, Yamada T, Murata K, Nakamura H, Ushida K. 2019. Role of coprophagy in the cecal microbiome development of an herbivorous bird Japanese rock ptarmigan. J Vet Med Sci 81:1389–1399.

## References

(1) Clermont O, Christenson JK, Denamur E, Gordon DM. 2013. The Clermont Escherichia coli phylo-typing method revisited: improvement of specificity and detection of new phylo-groups. Environ Microbiol Rep 5:58–65.

